# CCR6 is essential for effective immunity against *Mycobacterium tuberculosis* infection in mice

**DOI:** 10.1101/2025.08.07.669071

**Authors:** Summer J. Harris, Oyindamola O. Adefisayo, Rachel K. Meade, E. Ashley Moseman, Charlie J. Pyle, Clare M. Smith

## Abstract

Tuberculosis (TB) is a global epidemic that has threatened human health throughout recorded history. TB disease, caused by infection with *Mycobacterium tuberculosis (M.tb),* is heterogeneous between individuals, and clinical TB outcomes are impacted by genetic and environmental risk factors. Infected individuals must maintain a careful balance of cytokines and chemokines following *M.tb* infection to eliminate or contain the bacteria without causing excessive inflammation and lung damage. The CC chemokine receptor 6 (CCR6) is expressed by several immune cell populations that are classically involved in the pathogenesis of *M.tb* infection. However, the precise functions of CCR6 in the context of TB disease remain underexplored. Here, we show that mice lacking the CCR6 receptor (CCR6 KO) fail to restrict bacterial burden in the lungs, leading to increased disseminated infection at 4-weeks post-infection compared to wild-type C57BL/6J (B6) mice, with a more pronounced effect in females. CCR6 KO mice also developed necrotic pulmonary lesions, phenotypically distinct from B6 mice, and produced elevated levels of pro-inflammatory cytokines and chemokines at the onset of adaptive immunity, particularly IL-17. Long-term infection experiments revealed that the absence of CCR6 enhances risk for mortality following *M.tb* infection, particularly in females. This study provides insights into the role of CCR6 during TB pathogenesis and establishes its importance in maintaining protective immunity against *M.tb* within the context of a genetically tractable mouse model that forms necrotic pulmonary granulomas.

## Introduction

An estimated 10.8 million people worldwide were infected with tuberculosis (TB) in 2023, and among those cases, 1.25 million people succumbed to disease (1). 90% of people who become infected with *Mycobacterium tuberculosis* (*M.tb*), the causal agent of TB, successfully control infection, while the other 10% progress to an active disease state, usually as a result of a failure of the host’s immune responses (2). Effective host resistance to *M.tb* necessitates coordinated innate and adaptive immune responses (3). As such, numerous leukocyte effector cells involved in both the innate and adaptive response, such as neutrophils and T cells, are recruited to the site of infection where they play diverse roles necessary for bacterial containment and elimination (3, 4). Recruitment of these cell types is dependent on the immune milieu consisting of cytokines and chemokines (5–10). This is illustrated in a transcriptional profiling study of the innate immune response in mouse lung tissue following low-dose *M.tb* infection which revealed that many cytokines and chemokines are strongly upregulated 12 – 21 days following infection (11) and are likely correlated with the recruitment of immune cell populations to the site of infection. Cytokine- and chemokine-mediated recruitment and activation of microbicidal immune cells is crucial for controlling bacterial growth but requires balance with tolerogenic signaling to avoid causing excessive tissue injury (12), which may exacerbate disease outcomes in the host. In addition to direct microbicidal activity, recruited cells also participate in the formation of granulomas which are characterized as aggregates of immune cells around infecting bacterial cells (13). Chemokines play a particular role in granuloma formation (12, 14), which relies on a balance of signals to determine its function either as a protective niche for the bacteria or as a host-defense mechanism (15).

Despite their importance, elucidating the specific roles and impacts of cytokines and chemokines during infection remains challenging. This complexity arises from the fact that most cytokines and chemokines have multiple cellular targets and engage in intricate, redundant signaling pathways (12, 14). This makes the single receptor-single cytokine/chemokine pairs particularly attractive for study as they provide a scaffold for detangling intricate patterns of cytokine and chemokine responses during TB infection. One such chemokine-chemokine receptor pair is composed of CCL20 and CCR6. CCR6, which has no other known ligands (16), is expressed on immature dendritic cells (DC), innate lymphoid cells (ILCs), B cells, regulatory CD4 T cells (Tregs) and Th17 cells (17). The CCR6-CCL20 axis has been predominantly studied for its role in autoinflammatory conditions like inflammatory bowel disease (IBD) (17) and cancer (18) but has also been implicated in various infectious diseases such as human immunodeficiency virus (HIV) (19, 20), pneumococcal meningitis (21) and Helicobacter pylori gastritis (22, 23). Studies have shown that CCL20 is upregulated in *M.tb*-infected monocytes (24), activated peripheral blood mononuclear cells (PBMCs) from human patients with active TB (25), and in mice infected via intratracheal route with the *Mycobacterium bovis*-derived vaccine strain Bacille Calmette-Guérin (BCG) (26). In both non-human primates (NHPs) (27) and human patients (28) with active TB, the majority of airway CD4^+^ T cells co-expressed CCR6 and another chemokine receptor CXCR3 (27, 28). Although IFN-γ production is known to be essential for an effective host immune response to *M.tb* (29–31), Proulx and colleagues (32) characterized *M.tb*-resistant genetically diverse mouse strains that produce low levels of IFN-γ following infection, finding that these mice expressed higher levels of CCR6 on their CD4^+^ T cells correlating strongly with bacterial control. This is particularly relevant as CD4+ T cells can mediate mycobacterial control in both IFN-γ-dependent and IFN-γ-independent manners (33). Collectively, these data suggest that CCR6 is a prominent feature of the host adaptive immune response to *M.tb* infection that can operate independently of canonical IFN-γ-driven TB immunity and warrants further investigation.

To better understand the role of CCR6 during TB pathogenesis, we infected CCR6 receptor knockout (CCR6 KO) mice with aerosolized *M.tb* and compared disease phenotypes with co-infected B6 controls. In contrast to a previous study reporting that CCR6 is not required for the establishment of T cell-mediated adaptive immune responses or bacterial clearance following avirulent BCG infection (26), we show that the CCR6 deficiency disrupts adaptive immune control of virulent *M.tb*, resulting in a survival defect. Unlike in B6 mice that canonically fail to form organized *M.tb*-containing granulomas characteristic of human pulmonary TB (34), we identified multiple instances of organized necrotic lesions in infected CCR6 KO mice. These findings suggest that CCR6 signaling may play a role in shaping the pulmonary immune landscape in infected mice. Moreover, CCR6 KO mice may represent a novel and tractable system for investigating granuloma formation and dynamics following *M.tb* infection on a B6 background, enhancing our capacity to study granuloma biology *in vivo* and explore its impacts on TB outcome and on the pharmacodynamics of investigational therapies.

## Results

### CCR6 KO mice have elevated bacterial burden at 4-weeks post-infection

To investigate the role of CCR6 during *M.tb* infection *in vivo*, we infected CCR6 KO mice of both sexes via aerosol route and compared their infection dynamics to B6 mice. We observed no significant differences at 2- or 3-weeks post-infection, when the innate immune response is active (**Figure 1A**). However, at the onset of adaptive immunity around 4-weeks post-infection (35) CCR6 KO mice yielded significantly higher bacterial burden from their lungs (**Figure 1A**; p=0.0346) and spleens (**Figure 1B**; p=0.0284). After stratifying by sex, we observed a sex-specific bacterial burden defect in the lungs of CCR6 KO mice. Compared to B6 females, significantly higher bacterial burden was measured in the lungs of CCR6 KO females at 4-weeks post-infection (**Figure 1C**; p=0.0462). No significant lung burden difference was observed between CCR6 KO and B6 males (**Figure 1C**; p=0.373). The enhanced female burden defect was limited to the lung, as no sex-specific differences in bacterial control were observed in the spleen at 4-weeks post-infection (**Figure 1D**).

**Figure 1.**
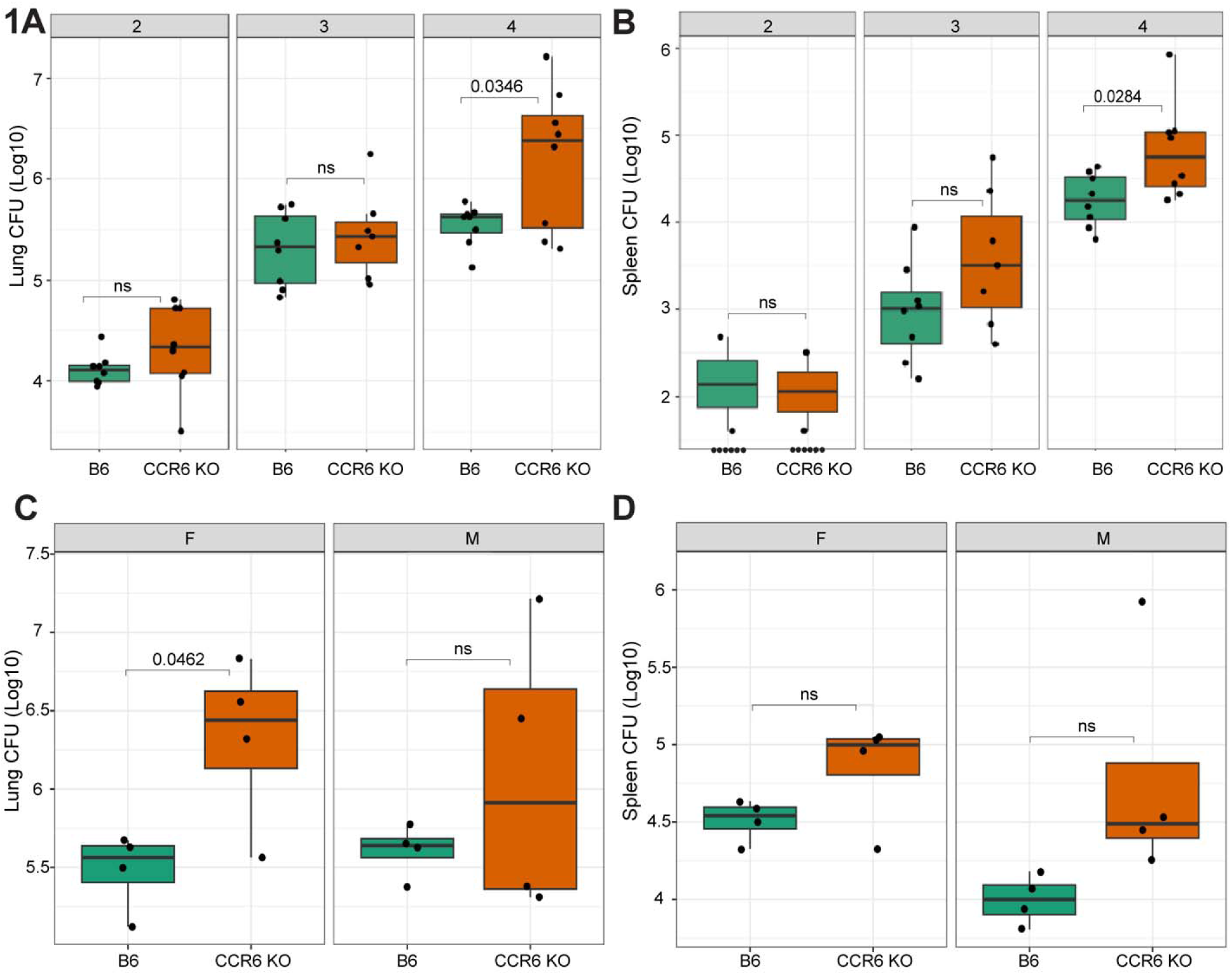
CCR6 KO mice have higher bacterial burden in the lung and spleen at 4-weeks post-infection. Bacterial burden measured from (**A**) lung and (**B**) spleen homogenate at 2-, 3-, and 4-weeks post-infection from CCR6 KO and B6 mice infected with *M.tb* H37Rv (n=8 mice per strain; both sexes included in equivalent quantity). Bacterial burden levels, stratified by sex, at 4-weeks post-infection from (**C**) lung and (**D**) spleen (n=4 mice per strain and sex). Hypothesis testing was performed by two-way ANOVA and Tukey’s *post hoc* test on log_10_ transformed values.

### CCR6 KO mice develop necrotic lung lesions

To explore the pulmonary immune landscape underlying this burden defect, we assessed the impact of CCR6 deficiency on infected lung pathology. We observed two major lesion phenotypes in the CCR6 KO lungs at 4-weeks post-infection. The first type was visibly similar to lesions observed in *M.tb*-infected B6 mice (**Figure 2A**, middle panel), which are characterized by disorganized clusters of primarily macrophages and lymphocytes (36). The second type of lesion was more organized, with a central necrotic core (**Figure 2A**, right panel). We next assessed lung damage using an artificial intelligence (AI) model (as described in the Methods section) to quantify cellular infiltration and necrosis within the lungs. Using that model, we found no significant differences in lesion size relative to the entire lung area between CCR6 KO mice and B6 controls (**Figure 2B**). We also did not observe a significant difference in areas of necrosis within the lesion between mouse lungs from CCR6 KO mice and B6 controls (**Figure 2C**). However, a subset of CCR6 KO mice developed pulmonary lesions with prominently expanded areas of necrosis (**Figure 2A** & **2C**). Histopathological analysis of lungs from independent cohorts of male CCR6 KO mice at 10-15 weeks post-infection, revealed that 30% (n= 3 of 10) of those animals formed well-organized, necrotic granulomas (**Figure 2D**, left panel). These granulomas resembled mycobacterial granulomas found in human patients (37), NHPs (38), and zebrafish (39), with a well-organized periphery comprised of immune cells, including epithelioid macrophages and foamy macrophages, surrounding a necrotic core (**Figure 2D**, right panel). Masson’s trichrome staining revealed that *M.tb* granulomas in CCR6 KO mice are surrounded by a collagen-rich fibrotic layer (**Figure 2E**), which is also present in mature human TB granulomas (40).

**Figure 2.**
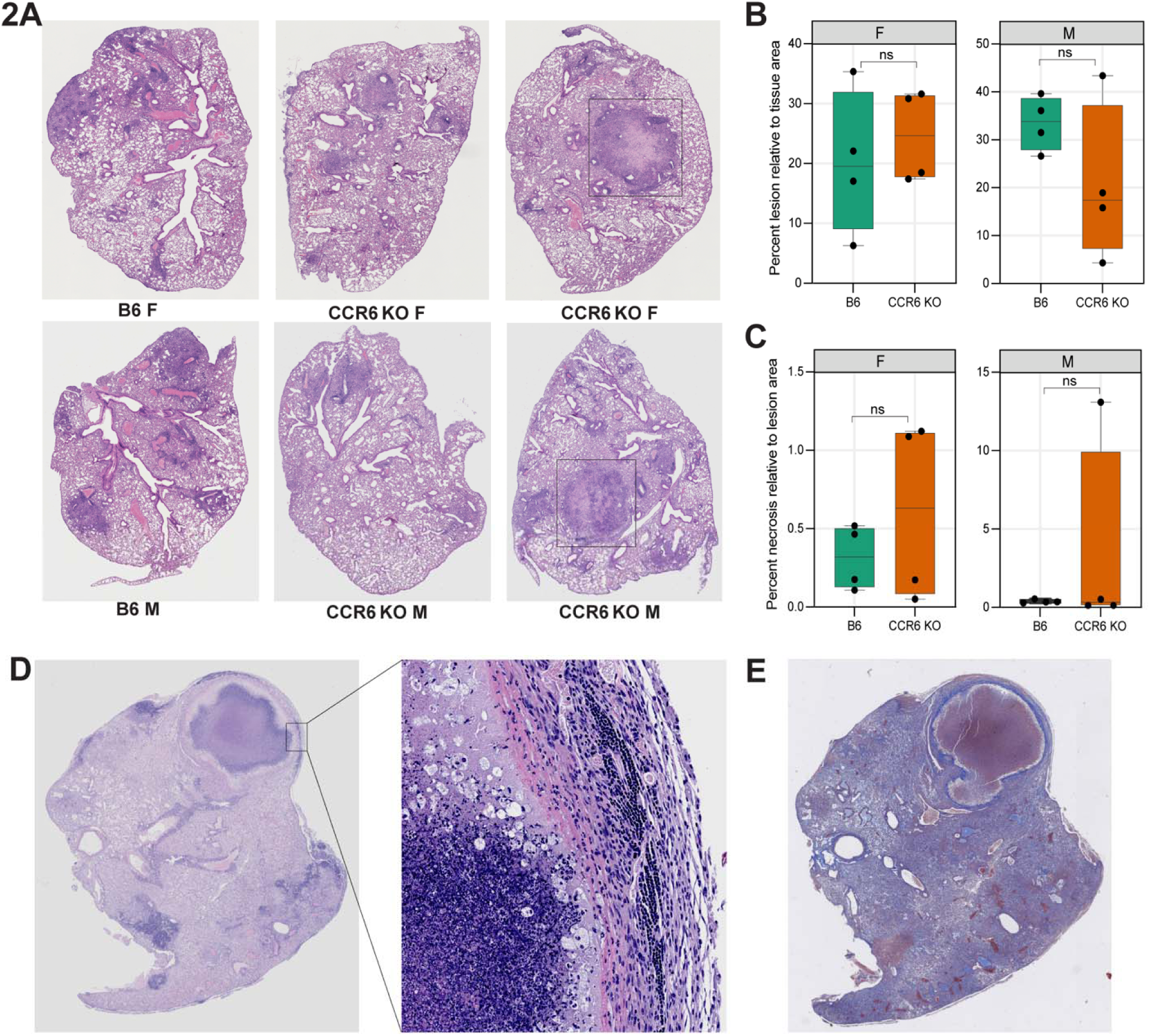
CCR6 KO mice develop necrotic lesions. (**A**) Representative whole-slide scans of H&E-stained lung sections from male and female CCR6 KO and B6 mice, collected at 4-weeks post-infection. Areas of organized lesion formation are indicated by a black box. Quantification of H&E-stained images demonstrating (**B**) the percent lesion area relative to total tissue area and (**C**) the percent necrotic area relative to the quantified lesion area, with each dot representing an individual mouse at 4-weeks post-infection. For panels **B** & **C**, no significant difference could be detected by unpaired *t*-test with Welch’s correction. (**D**) Representative H&E-stained whole-lung section from a male CCR6 KO mice at 15 weeks post-infection (left panel). The right panel is a 40X magnification of the boxed area from the left panel showing the periphery of a well-organized, necrotic lesion. (**E**) Masson’s trichrome-stained slide from the same representative sample in panel **D**, indicating the fibrotic capsule (deep blue stain) surrounding the lesion.

### *M.tb*-infected CCR6 KO mice have elevated IL-17 levels at 4-weeks post-infection

We next evaluated the impact of CCR6 deficiency on pulmonary inflammation in *M.tb*-infected mice. We conducted immunological profiling on lung homogenate from *M.tb*-infected CCR6 KO mice and B6 controls via multiplexed ELISA. At 4-weeks post-infection, CCR6 KO mice produced elevated levels of numerous cytokines and chemokines including inflammatory cytokines IL-2 and IL-17, neutrophil/ macrophage maturation cytokines M-CSF, G-CSF and GM-CSF, as well as the neutrophil attractant chemokines KC (CXCL1), LIX (CXCL5), MIP-1α, MIP-1β, and MIP-2 (CXCL2) (**Figure 3A**). Conversely, CCR6 KO mice have reduced levels of angiogenesis factor VEGF which has been shown to have CCR6 dependent expression in cancer (41, 42). We sought to identify unique features of the CCR6 KO inflammatory signature. We conducted sparse Partial Least Squares Discriminant Analysis (sPLS-DA) across cytokine and chemokine levels as well as lung and spleen burden measurements from CCR6 KO and B6 lung samples at 4-weeks post-infection (**Figure 3B**). IL-17 and LIX were identified as the strongest features underlying sparse component 1, which explained 51% of the variance between groups (**Figure 3C**). Significant differences in IL-17 and LIX by host genotype were only observed at the 4-week timepoint (**Figure 3D**) and were not impacted by sex.

**Figure 3.**
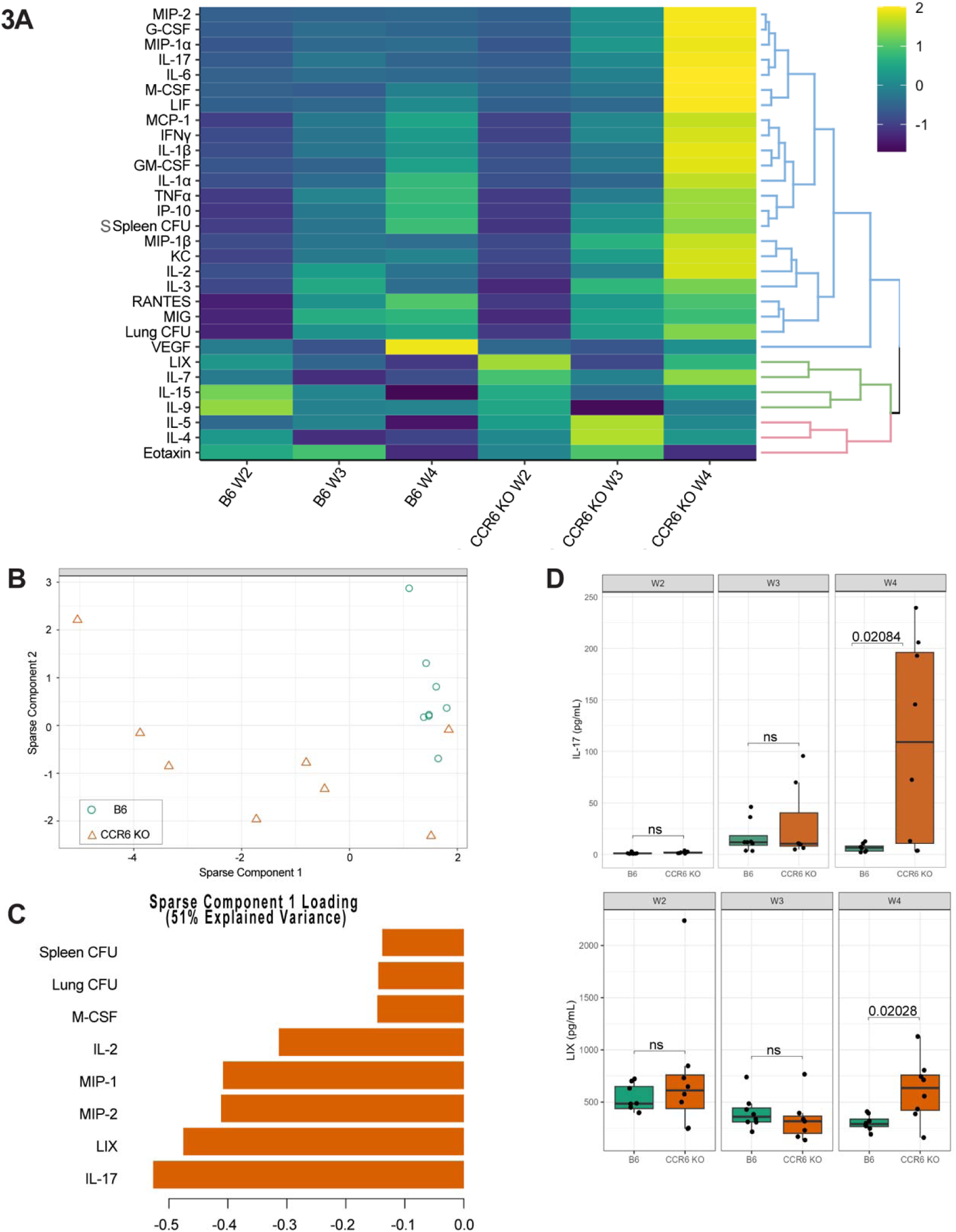
IL-17 and LIX levels dominate the immune responses in the lungs of *M.tb*-infected CCR6 KO and B6 mice. (**A**) Heatmap depicting scaled phenotypes at 2-weeks (W2), 3-weeks (W3), and 4-weeks (W4) post-infection, hierarchically clustered and divided into 3 k clusters. (**B**) Individual mice plotted against the first two sPLS-DA components which explained the greatest variance in the data from measurements collected 4-weeks post-infection. (**C**) Phenotype loadings contributing to the first sparse component with a 51% explained variance. (**D**) IL-17 and LIX levels measured from lung homogenate by multiplex ELISA at 2-, 3- and 4-weeks post-infection. Hypothesis testing was performed by two-way ANOVA and Tukey’s *post hoc* test.

### *M.tb* susceptibility in CCR6 KO mice is sex dependent

To understand the long-term impact of the observed defects in adaptive immune control of *M.tb*, we investigated post-infection survival in CCR6 KO mice. Prompted by our observations of elevated bacterial burden and inflammatory cytokine and chemokine levels in CCR6 KO lungs at 4-weeks post-infection, we investigated the impact of CCR6 ablation on long-term *M.tb* infection outcomes. We infected independent cohorts of CCR6 KO and B6 control mice via aerosol route and evaluated clinical indicators of TB progression over a 70-week period. IFNGR KO mice, which are canonically susceptible to *M.tb* infection (43), were included as controls to confirm successful aerosol infection. Over the study period, animals were weighed bi-weekly to monitor for signs of severe disease and were euthanized at humane endpoints in accordance with IACUC-approved protocols. Compared to B6 controls, infected CCR6 KO mice lost a greater proportion of their initial body weight on average and did not recover within the observation period (**Figure 4A**). This wasting phenotype was more pronounced in female CCR6 KO mice after 30 weeks of infection (**Figure 4B**). Conversely, male CCR6 KO mice did not exhibit severe wasting during infection, with the mean change in body weight steadily increasing throughout the observation period. CCR6 ablation was associated with a significant overall reduction in survival (**Figure 4C**; p=0.03). Consistent with observed differences in wasting throughout infection, only female CCR6 KO mice were at significantly increased risk of mortality following *M.tb* infection (**Figure 4D**, p=6e^-4^ in females, p=0.3 in males). Collectively, these data suggest that functional CCR6 is essential to supporting host resistance to *M.tb* infection, particularly in females.

**Figure 4.**
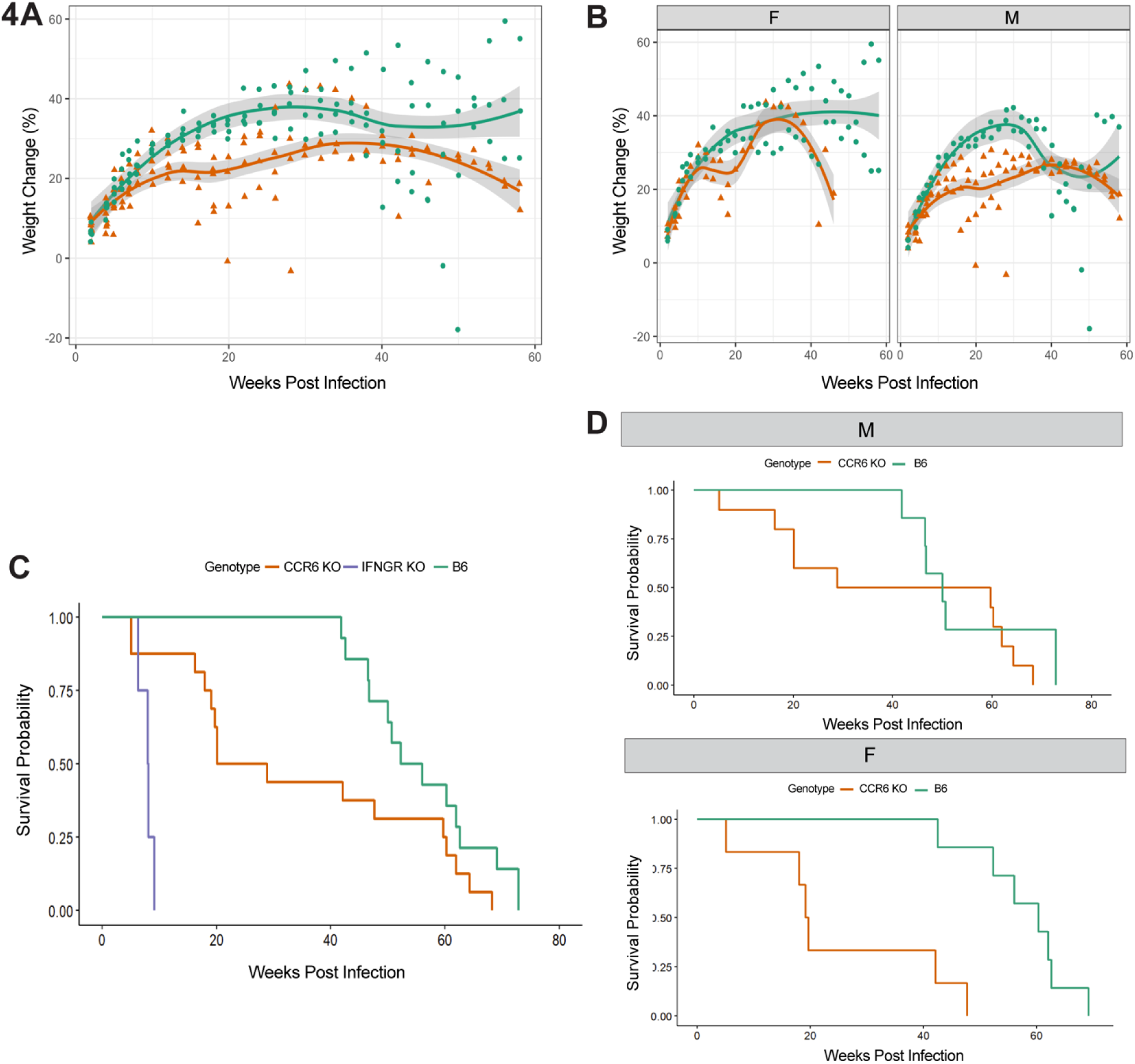
CCR6 KO mice exhibit increased *M.tb* susceptibility and higher post-infection mortality risk. Mice were challenged with *M.tb* H37Rv and weighed bi-weekly for 70 weeks to monitor disease progression. Orange lines represent the average weight of CCR6 KO mice, and green lines represent the average weight of B6 mice (**A**) Kinetics of percent weight change compared to initial body weight at day 0 for the complete cohort. (**B**) highlighting the kinetics of percent weight change from initial body weight stratified by sex (n=7-10 mice per genotype and sex). (**C**) Kaplan-Meier survival estimates of *M.tb*-infected CCR6 KO, IFNGR KO, and B6 for the complete cohort. (**D**) Kaplan-Meier survival estimates stratified by sex for *M.tb*-infected CCR6 KO and B6 mice.

## Discussion

Research on latent tuberculosis infection (LTBI) in humans (44) and NHPs (27) indicates that the majority of TB specific CD4+ T cells also express CCR6. Nevertheless, the role of CCR6 during virulent mycobacterial infection remain undetermined. Using a mouse model with whole-animal CCR6 ablation, we have determined that CCR6 is necessary for restriction of bacterial burden, likely primarily through promotion of an effective adaptive immune response. We show that CCR6 ablation impacts bacterial burden after adaptive immunity onset at 4-weeks post infection, particularly in female mice. In addition to increased bacterial burden in CCR6 KO mice, we observed necrotic lesions starting at 4-weeks and progressing to fully encapsulated necrotic human-like granulomas by 15-weeks. Immune profiling of infected lungs showed elevated cytokine and chemokine expression in CCR6 KO mice, with high IL-17 as a strong feature of TB infection in the absence of CCR6. Finally, we show increased mortality risk in female CCR6 KO mice, correlating with increased bacterial burden.

Genetic variation in human *CCR6* has been shown to mediate disease severity across several inflammatory diseases. Genome-wide association studies in cohorts with rheumatoid arthritis, primary biliary cholangitis, and several other inflammatory conditions have identified strong associations between *CCR6* polymorphisms and disease severity (45–47). In agreement with these studies, we report that CCR6 ablation *in vivo* meaningfully enhances TB severity, especially in females.

Based on our results highlighting elevated IL-17 levels in CCR6 KO mice at 4-weeks post-infection and the known roles of IL-17 in supporting the adaptive immune response to mycobacterial infection (48–50) we hypothesize that the impact of CCR6 signaling on the adaptive immune response is associated with its promotion of the Th17 response. It has been previously demonstrated that IL-17-producing CD4+ T cells constitute the predominant CCR6-expressing population in both experimentally induced Th17 states and in an experimental animal model of autoimmune disease (51, 52). CD4^+^ T cells, with elevated CCR6 expression migrate to the lungs during *M.tb* infection in humans (28, 44), NHPs (27), and mice (32). Although CD4^+^ Th17 cells are known producers of IL-17, Lockhart and colleagues (53) identified gamma-delta (γ/δ) T cells as the main producers of IL-17 during murine TB. Importantly, previous studies in CCR6 KO mice have shown that CCR6 ablation does not impact γ/δ T cell recruitment to the lungs following BCG airway infection (26). In concordance with this previous study of BCG-infected CCR6 KO mice, we found that elevated IL-17 expression was a particular hallmark of mycobacterial infection in CCR6 KO mice (26). Elevated IL-17 levels in CCR6 KO mice suggest adaptive immune dysregulation when CCR6 signaling is absent, likely resulting in the observed disease susceptibility to *M.tb* infection *in vivo*.

IL-17 facilitates immune protection by enhancing neutrophil recruitment to the site of infection (54, 55). However, it is well established that maintaining balanced neutrophil levels is crucial during chronic TB. In mice, IL-17-mediated neutrophilia is associated with chronic lung inflammation and increased mortality (56, 57). We found that CCR6 KO mice exhibit increased mortality rates, which is particularly pronounced in female mice. This observation highlights a disparity in the commonly noted higher TB mortality rates observed in male mice (58, 59). Although we did not observe a significant increase in neutrophil infiltration in the lungs of CCR6 KO mice compared to B6 controls, we did identify the presence of organized lesions in the lungs of CCR6 KO mice as early as 4-weeks post-infection. Future investigations into the immune cell profiles of CCR6 KO mice at subsequent time points are essential for elucidating alterations in immune cell trafficking to the lungs and the occurrence of neutrophilia. Such analyses will provide valuable insights into the formation of necrotic granulomas.

Necrotic granuloma formation is a hallmark of chronic TB disease in humans and has been studied extensively as a determinant of pathogenesis using a variety of animal models, including NHPs and zebrafish (15, 39, 60, 61). Classical necrotic lesions are largely absent in B6 and other common laboratory mouse strains during *M.tb* infection (62, 63). Although our observation of necrotic granulomas in *M.tb*-infected CCR6 KO mice is not unprecedented in the context of murine TB models (64–66), it offers a compelling new model for investigating the role of granulomas in TB pathogenesis. While the precise role of CCR6 in necrotic granuloma formation remains to mechanistically dissected, this study provides an important step in understanding the complex immunological signaling that coordinates the granulomatous and adaptive immune response to infection with *M.tb*.

## Materials and Methods

### Ethics Statement

Mouse studies were approved by the Institutional Animal Care and Use Committee (IACUC) at Duke University (Protocol numbers: A204-23-10 and A221-20-11), using the recommendations from the Guide for the Care and Use of Laboratory Animals of the National Institute of Health and the Office of Laboratory Animal Welfare.

### Mouse strains and infection with *M.tb*

C57BL/6J (#000664), B6.129P2-*Ccr6^tm1Dgen^*/J (CCR6 KO; #005793), and B6.129S7-*Ifngr1^tm1Agt^*/J (IFNGR KO; #003288) mice were purchased from The Jackson Laboratory. CCR6 KO mice, which are maintained on a C57BL/6J (B6) genetic background, were subsequently bred at Duke. All mice were housed in a specific pathogen-free facility under standard conditions (12 h light/dark, food and water *ad libitum*). Male and female mice were infected between 8 and 12 weeks of age with *M.tb* H37Rv. *M.tb* strains used for *in vivo* infection were verified to be positive for the essential virulence factor, phthiocerol dimycocerosate (PDIM). *M.tb* was cultured in 7H9 media supplemented with 10% oleic acid-albumin-dextrose-catalase (OADC) enrichment (Middlebrook), 0.2% glycerol, and 0.05% Tyloxapol (Fisher). Prior to infections, *M.tb* cultures were washed and resuspended in phosphate-buffered saline containing 0.05% Tween 80 (PBS-T). Bacterial aggregates were then broken into single cells using a blunt needle before diluting to desired concentration for infection. An inoculum between 90 and 200 colony-forming units (CFU) was delivered by an aerosol-generating Glas-col chamber. To determine the inoculation dose, groups of 4 mice were euthanized at 24 hours post-infection, and CFU were enumerated from lung homogenates, as described below.

### Bacterial burden quantification

To quantify CFU, mice were euthanized in accordance with IACUC-approved endpoints by overdose of isoflurane (Covetrus), and the lungs and spleens were aseptically removed. For enumeration of viable bacteria, organs were individually homogenized in PBS-T by bead beating (MP Biomedical), and 10-fold dilutions were plated on 7H10 agar (Middlebrook) plates containing 10% OADC enrichment (Middlebrook), and 50 µg/mL Carbenicillin, 10 µg/mL Amphotericin B, 25 µg/mL Polymyxin B, and 20 µg/mL Trimethoprim (Sigma). Plates were incubated at 37°C for 21 days, and individual *M.tb* colonies were enumerated to calculate CFU.

### Quantification of cytokines in tissue homogenate

Following homogenization, infected lung samples were centrifuged through 0.2µM filters to remove cellular debris. Thirty-two cytokines and chemokines were quantified via Discovery Assay (Eve Technology; MD31). In this assay, fluorescence intensity values were measured for all samples alongside a standard curve to estimate the expected concentrations for each cytokine. Observed cytokine concentration values (reported in pg/mL) were then interpolated for each sample and cytokine using the standard curve and fluorescence intensity values. From this panel, IL-10, IL-12p40, IL-12p70, and IL-13 fell below the detectable limit for all samples and were excluded from further analysis.

### Lung pathology and estimation of inflammation

At 2-, 3- and 4-weeks post-infection one lung lobe from each mouse was collected for histology. Each lobe was fixed in 10% neutral-buffered formalin (VWR) for 24-48 hours then washed 3 times with 70% ethanol. Lungs were submitted to the Duke BioRepository and Precision Pathology Center (BRPC) core where they were paraffin-embedded, sectioned at 5µM, and stained with hematoxylin and eosin (H&E). Each whole-lung section was scanned at either 10X or 40X using an Echo Revolution microscope. Whole-slide images (at 40X magnification) were analyzed using Aiforia® Create (Version 6.2; Aiforia Technologies Plc, Helsinki, Finland), a cloud-based artificial intelligence (AI) platform designed for training, deploying, and validating pathology image analysis models(67–69).A total of 153 whole slide images of mouse lung tissue were used for the AI model development. Out of the 153 images, 133 were used for the model training, and 20 images were left out of the training set for AI model validation purposes. Validation was performed according to a three-point grading system. Four individual validators gave scores for the AI model analysis results according to how close in their opinion the AI model captured the relevant features. The analytical scheme is represented in **Supplemental Figure 1**. For preparation of whole-lung histology samples from long-term experiments, lungs were first perfused with 3mL of 10% neutral-buffered formalin through the trachea using a 22-gauge needle before being placed in 10% neutral-buffered formalin for 48 hours and processed as described above.

### Statistical Analysis

Hypothesis testing and data visualization were performed using R statistical software (version 4.3.1). Hypothesis testing for differences in bacterial burden, lung damage, and cytokine levels by host genotype was performed by two-way ANOVA and Tukey’s *post hoc* test. Differences in survival by host genotype were estimated using Kaplan-Meier analysis in the R package survminer, and statistical significance was assessed via Mantel-Cox hypothesis testing.

## Acknowledgements

The authors acknowledge Joshua Tolliver, Kaley M. Wilburn, Alwyn M. V. Ecker, and Marco T. P. Gontijo for technical assistance. This work was funded by an NIH Director’s New Innovator Award AI183152 (C.M.S) and a Pew Scholars award (C.M.S). Biocontainment work performed in the Duke Regional Biocontainment Laboratory received partial support for construction and renovation from NIAID (UC6-AI058607 and G20-AI167200) and facility support from the NIH (UC7-AI180254). The sponsors or funders did not play any role in the study design, data collection and analysis, decision to publish, or preparation of the manuscript.

## Author Contributions

S.J. Harris: conceptualization, data curation, formal analysis, investigation, validation, visualization, and writing-original draft, review and editing. O.O. Adefisayo: data curation, formal analysis, investigation, validation, visualization, and writing-original draft, review and editing. R.K. Meade: data curation, formal analysis, investigation and writing-review and editing. E.A. Moseman: Resources and writing-review and editing. C.J. Pyle: investigation, validation, visualization, and writing-original draft, review and editing. C.M. Smith: conceptualization, funding acquisition, project administration, supervision, and writing-original draft, review and editing.

