## Supplemental Figure 1 for "CCR6 is essential for effective immunity against *Mycobacterium tuberculosis* infection in mice"

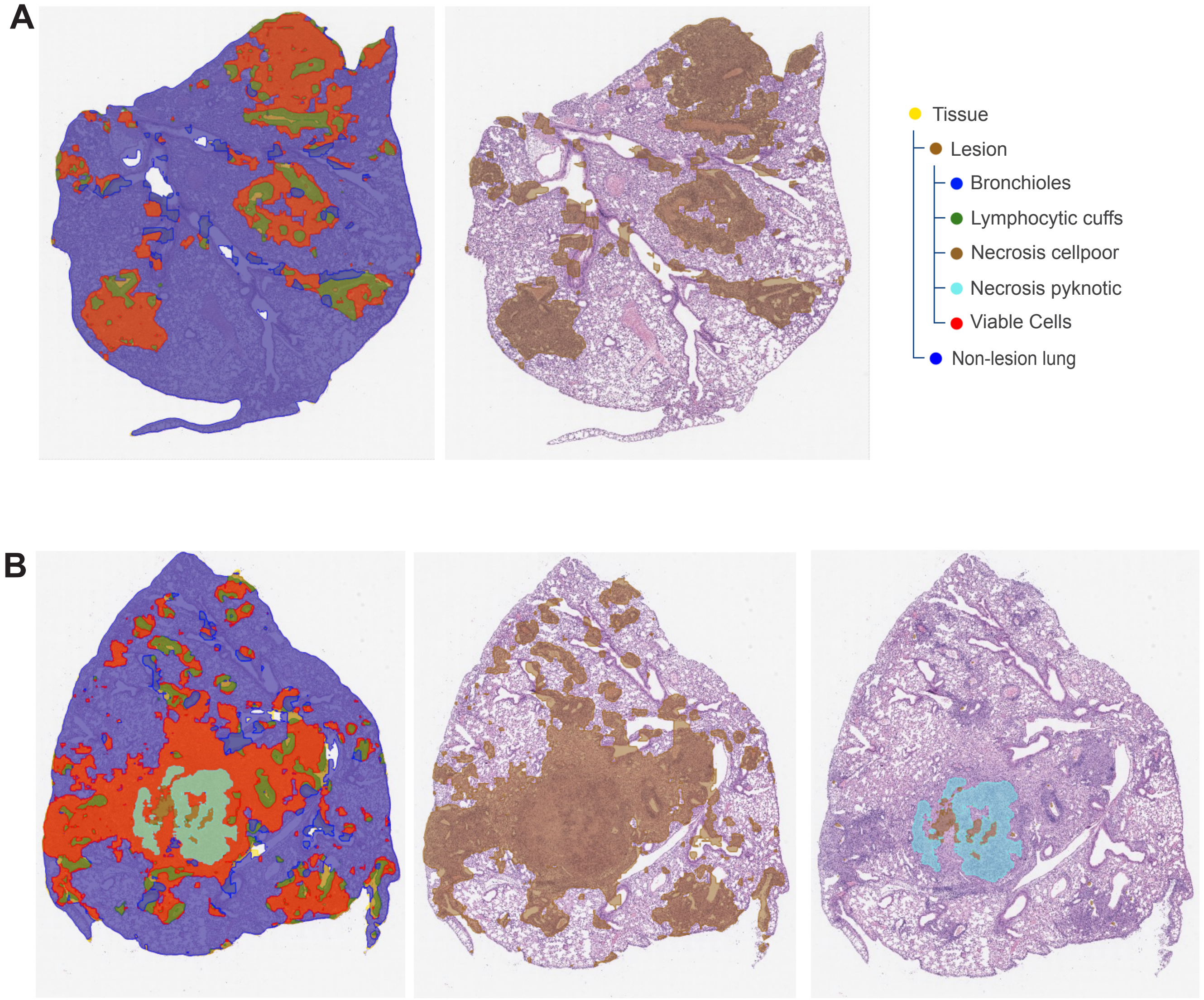


**Supplemental Figure 1. Cellular infiltration and necrosis in the lung.** An AI-based model analyzed whole-lung scans of H&E-stained sections from both (A) B6 and (B) CCR6 KO male mice. For each image, 3 separate layers were analyzed: tissue vs. background, lesion vs. non-lesion (damage vs. healthy tissue), and lesion sub-regions (bronchioles, lymphocytic cuffs, necrosis cellpoor, necrosis pyknotic, and viable cells). In both A & B the left panel depicts all 3 layers overlapped. The middle panel shows the lesion vs. non-lesion layer. The right panel in A shows the color assignment for each layer and sub-region. The right panel in B only represents the necrosis cellpoor (brown) and necrosis pyknotic (blue) subregions in the lesion layer in CCR6 KO mice, which was not detected in B6 controls.
